# Computation of history-dependent mechanical damage of axonal fiber tracts in the brain: towards tracking sub-concussive and occupational damage to the brain

**DOI:** 10.1101/346700

**Authors:** Jesse I. Gerber, Harsha T. Garimella, Reuben H. Kraft

## Abstract

Finite element models are frequently used to simulate traumatic brain injuries. However, current models are unable to capture the progressive damage caused by repeated head trauma. In this work, we propose a method for computing the history-dependent mechanical damage of axonal fiber bundle tracts in the brain. Through the introduction of multiple damage models, we provide the ability to link consecutive head impact simulations, so that potential injury to the brain can be tracked over time. In addition, internal damage variables are used to degrade the mechanical response of each axonal fiber bundle element. As a result, the stiffness of the aggregate tissue decreases as damage evolves. To counteract this degenerative process, we have also introduced a preliminary healing model that reverses the accumulated damage, based on a user-specified healing duration. Using two detailed examples, we demonstrate that damage produces a significant decrease in fiber stress, which ultimately propagates to the tissue level and produces a measurable decrease in overall stiffness. These results suggest that damage modeling has the potential to enhance current brain simulation techniques and lead to new insights, especially in the study of repetitive head injuries.

## 1 INTRODUCTION

Traumatic brain injury (TBI) is a significant cause of death and long-term disability [1]. In the United States, there were 2.8 million TBI-related emergency department visits, hospitalizations, and deaths in 2013 [2]. The structure of the brain can be divided into the gray and white matter regions. In general, gray matter tissue forms the outer layer of the brain and contains the neuron cell bodies, while white matter tissue forms the central region of the brain and contains neuron cell extensions, known as axons. These tightly bundled axons form a dense communication network that transmits signals between the neuron cell bodies. While axons account for the majority of white matter tissue, they are surrounded by a mixture of other components, such as glial cells and the extracellular matrix (ECM). However, this surrounding host material is much softer than the axons [3] [4]. As a result, white matter tissue is commonly treated as a fiber-reinforced composite in mechanical simulations [5] [6] [7] [8]. During severe head impacts, brain deformations can result in the widespread stretching and shearing of axons. This kind of TBI is classified as diffuse axonal injury (DAI) and is typically associated with motor vehicle crashes, sports injuries, and military blast trauma [9] [10]. DAI is the primary injury mechanism in more than 40% of all hospitalized TBIs [11] and can result in both physical and cognitive impairments, which may be temporary or permanent [12].

In an effort to design safer products and study the underlying mechanisms of brain injury, there has been a significant push to develop biofidelic finite element (FE) models of the human head. In the future, this technology could be used to create a brain health monitoring system for humans functioning in extreme environments. Brain injuries, especially mild TBIs (mTBI), are difficult to diagnose with medical imaging [13] [14]. Therefore, when combined with wearable sensors, simulations could provide real-time diagnostics that could be used to screen for biomechanical injury and allow for early medical intervention [15]. At this time, a number of FE brain studies have specifically focused on the prediction of DAI [16] [17] [18] [19] [20]. Using a variety of methods, these models predict axonal strains, which are then compared with functional and mechanical tissue thresholds [21]. While these studies are extremely useful, they have been restricted to simulating separate isolated events and, to our knowledge, all head and neck FE models have used elastic constitutive models. At this time, there is no way to account for the accumulated damage that is caused by multiple head impacts. In this paper, we propose a computational method that provides a first step towards providing this capability.

We begin with the observation that soft biological tissues have been shown to degrade with successive loadings. For instance, tendons [22], intervertebral discs [23], and bioprosthetic heart valves [24] all exhibit stiffness degradation and finite life, when subjected to cyclic loading. In addition, brain tissue has been shown to experience significant stress softening [25] [26], also called the Mullins effect [27], after the initial loading cycle. This behavior has been successfully modeled for brain tissue [25] using the pseudo-elasticity theory [28], which we adopt in this paper. In this theory, the degree of softening is related to the maximum strain energy achieved throughout the previous loading history. Stress softening is a damage-induced inelastic phenomenon caused by structural rearrangement in the material. For soft biological tissues, the Mullins effect is not associated with fiber/matrix bond rupture and complete damage [29].

In the pseudo-elasticity theory, the degree of softening will only increase when the current strain energy exceeds its previous all-time maximum. Therefore, a cyclic load at constant strain amplitude would only cause damage on the first cycle. In order to address this shortcoming, we have also included a fatigue-induced stress softening model, which has recently been used to simulate fatigue damage in bioprosthetic heart valves [30]. At this time, the functional and mechanical fatigue properties of brain tissue have not yet been experimentally studied. However, based on the prevalence of fatigue damage in other soft biological tissues, we assume that brain tissue also experiences cycle-dependent degradation.

According to quasi-static tensile tests [25], the stress response of brain tissue gradually increases to a peak value and then decreases until final tissue separation, which occurs at some nonzero stress value. The softening that occurs before failure is typical of damaging fibrous materials and is caused by the progressive rupture of the filamentary structure [25]. The pseudo-elasticity theory and fatigue-induced stress softening models are not capable of degrading the stress response during the initial load cycle. Therefore, we implemented an additional fiber rupture damage parameter, in order to capture the observed post-yield stress softening behavior. Lastly, we implemented an additional matrix damage parameter, which is based the mean total damage of the adjacent fibers. As a result, we are able to approximate the full experimental stress response.

In line with our previous work [18], we use the embedded element method to incorporate the axonal fiber tracts in our FE model of white matter tissue. As a result, we are able to monitor the response of each fiber track segment and maintain a record of the associated damage parameters. When a simulation is complete, we export the final values of the damage parameters and then use them as the initial conditions for a future simulation. In this sense, we are able to bridge the gap between successive head impacts. In order to account for the healing process, we have provided a mechanism to partially or completely reverse the accumulated damage in each fiber. To activate this feature, a healing duration is specified when the future simulation is initialized. Healing has been modeled in various soft biological tissues, such as tendons [31], ligaments [32], and intervertebral discs [33]. However, due to the lack of experimental data, a constant healing rate is typically assumed. Similarly, we also assume a constant healing rate.

In this study, we present the details of the proposed stress softening, fatigue damage, fiber rupture damage, and healing models. Then, using example simulations, we demonstrate the features and mechanics of the overall computational framework. Finally, we conclude with a summary of behavior trends, current limitations, and future challenges.

## 2 METHODS

### 2.1 EMBEDDED ELEMENT METHOD

In this paper, we use our own custom explicit nonlinear FE code, which is based on the total Lagrangian formulation and central difference method [34]. The source code is available on GitHub (https://github.com/PSUCompBio/compbio), along with the input files and results associated with this study. A fiber-reinforced composite can be divided into separate matrix and fiber components, as illustrated in Fig. 1. The mechanical properties of the composite will depend on the properties of the constituents, the geometry of the fibers, and the distribution of the fibers within the matrix [35]. In white matter tissue, the mixture of cells surrounding the axons represents the matrix component and is modeled using 8-node trilinear hexahedral elements. The axonal fiber track segments are modeled using 2-node linear truss elements, which only deform by axial stretch. In addition, the fibers are assumed to be incompressible, which is the standard nonlinear approach in commercial software [36].

**Fig. 1.**
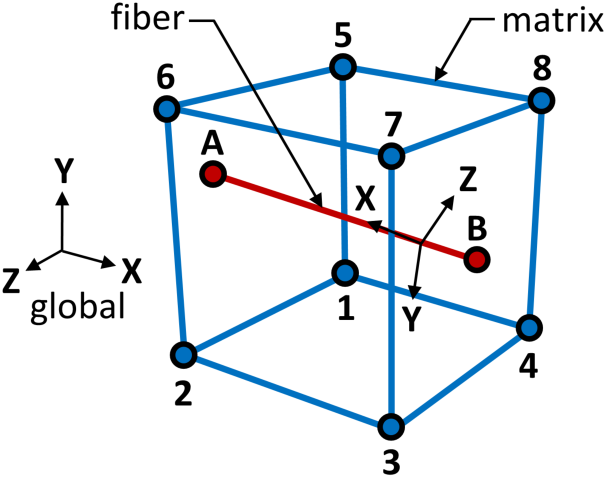
Simplified model of a composite material.

In our code, the unknowns are only calculated at the nodes of the matrix mesh. The fiber elements are incorporated using the embedded element method, where the core assumption is that the fiber nodes are bonded to the associated material points in the matrix element domain. We begin by using the standard solution algorithm [34] to calculate the displacements of the matrix element nodes. Next, we use these displacements, along with the matrix shape functions, to determine the displacements of the fiber element nodes. Then, we use these displacements to calculate the current stretch in each fiber element, which is then used, with the prescribed constitutive relation, to calculate the resulting uniaxial stress along each fiber axis.

In the standard solution algorithm [34], the primary task is to determine the internal forces at the nodes of the mesh. In a typical mesh, without embedded truss elements, the internal nodal forces are calculated as follows:

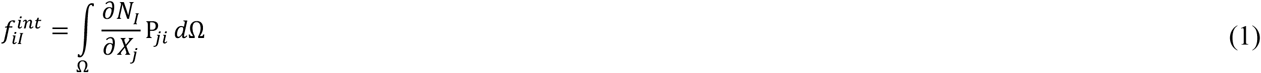

where lower case subscripts are used for components, upper case subscripts are used for nodal values, *N_I_* are the element shape functions, *X_J_* are the initial positions, and *P_ji_* are the components of the first Piola-Kirchhoff (PK1) stress tensor **P**. Now, in order to include the stiffening effect of the fiber elements, we modify Eq. (1) by including an additional integral over each fiber element domain. Therefore,

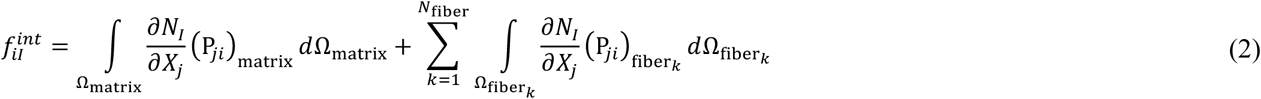

where *N_I_* are the matrix element shape functions in all terms. Note that, during the preprocessing stage, all fiber elements are split at the matrix element boundaries. Therefore, each fiber element is only associated with a single host matrix element.

### 2.2 MATERIAL MODELS

The following material models are based on our previous work [18]. As a result, the matrix material is modeled using the modified Mooney-Rivlin hyperelastic strain energy function [36], which is written as:

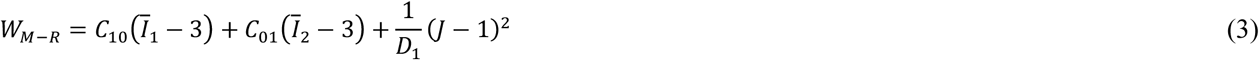

where 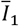 and 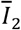 are the first and second deviatoric principal invariants of 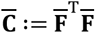, where 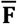 is the deviatoric part of **F**; *C*_10_, *C*_01_, and *D*_l_ are material parameters; and *J* ≔ det (**F**). The deviatoric principal invariants of 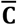 are related to the principal invariants of **C** ≔ **F**^T^**F** by:

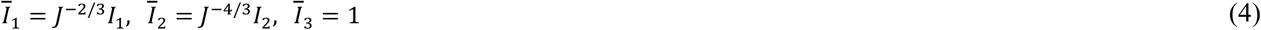

In order to provide a time-dependent response, we have also implemented a three-dimensional finite strain viscoelasticity model [37] for the matrix material, which is based on the generalized Maxwell-element. For brain tissue, viscoelastic behavior is typically associated with the isochoric part of deformation [38]. Therefore, the volumetric part of deformation is treated as purely elastic. The deviatoric stress in the reference configuration is calculated using the following recurrence relation:

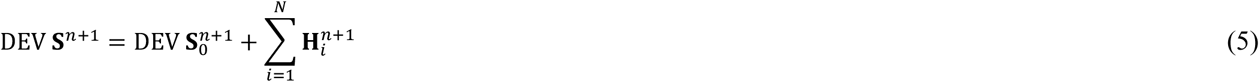

where DEV **S***^n^*^+l^ is the current deviatoric component of the second Piola-Kirchhoff (PK2) stress tensor, 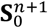 is the current deviatoric component of the elastic PK2 stress tensor, 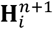 are the current internal stress variables, *n* + 1 is the current time, and *N* is a finite number of generalized Maxwell-elements. In addition,

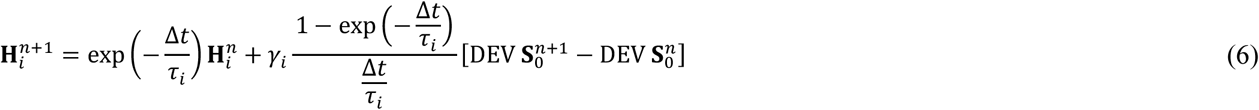

where Δ*t* = *t_n_*_+l_ − *t_n_* and *τ_i_* and *γ_i_* are scalar material parameters. The implemented viscoelasticity formulation [37] is very similar to the standard approach used in commercial software [36].

The fiber material is modeled using the modified Ogden hyperelastic strain energy function [36], which is written as:

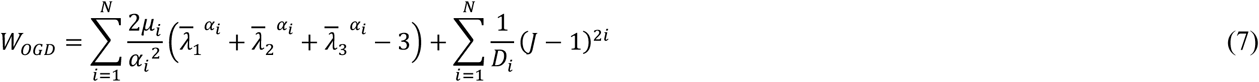

where 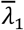, 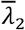, and 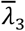 are the deviatoric principal stretches and *α_i_*, *μ_i_* and *D_i_* are material parameters. The deviatoric principal stretches are related to the principal stretches by:

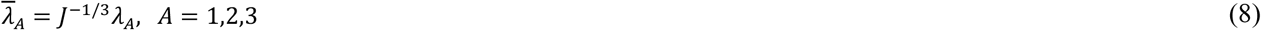

As mentioned earlier, the fibers are assumed to be incompressible. Also, as in our previous work [18], we select *N* = 1. Therefore, we now rewrite Eq. (7) as:

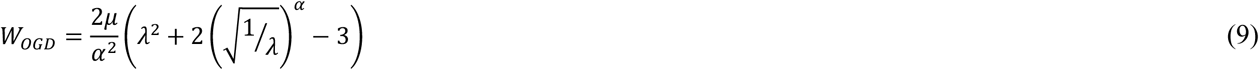

where *λ* is the axial stretch. Finally, we use Eq. (9) to express the uniaxial Cauchy stress as:

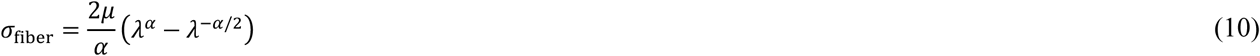

The material properties used in this study are listed in Table 1. The viscoelastic properties are identical to those used in our previous work [18]. The Mooney-Rivlin and Ogden properties are based on our previous work [18]. However, the values have been manually adjusted to approximate the experimental response [25] of brain tissue, as shown in Fig. 2b. To allow for rapid trial and error, we used a simplified FE model to select the material properties. The simplified FE model in shown in Fig. 2a and the final FE model is shown in Fig. 8c. The simplified FE model geometry was selected to approximately match the average fiber volume fraction of the final FE model.

**Table 1.** Material properties used for FE model.

| Mooney-Rivlin (matrix) | viscoelasticity (matrix) | Ogden (fiber) |
| --- | --- | --- |
| $\rho = 1040 \text{ kg/m}^3$ | $\gamma_1 = 1.977$ | $\rho = 1040 \text{ kg/m}^3$ |
| $C_{10} = -100 \text{ Pa}$ | $\tau_1 = 0.006694 \text{ sec}$ | $\alpha = 4.5$ |
| $C_{01} = 1.2 \text{ kPa}$ | $\gamma_2 = 0.0450$ | $\mu = 2.5 \text{ kPa}$ |
| $D_1 = 0.9091 \times 10^{-9} \text{ Pa}^{-1}$ | $\tau_2 = 0.1564 \text{ sec}$ | $D_1 = 0.9091 \times 10^{-9} \text{ Pa}^{-1}$ |

**Fig. 2.**
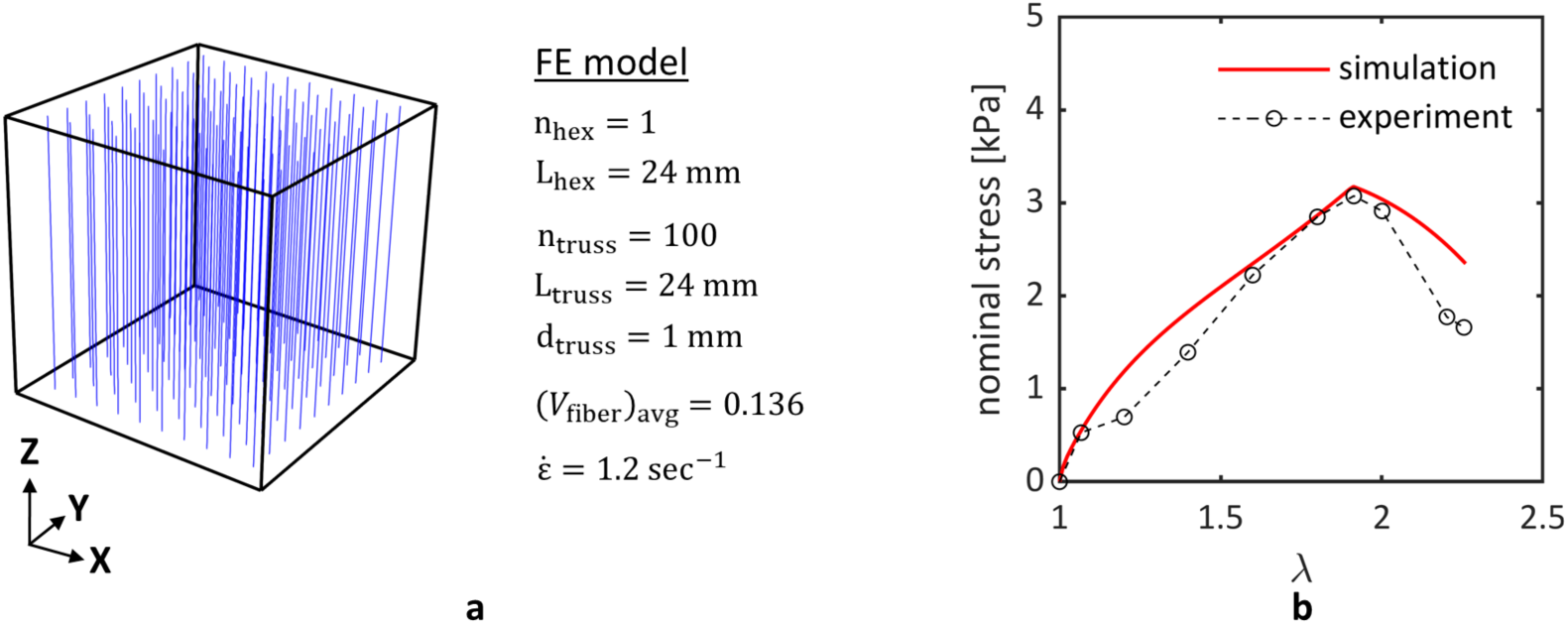
The (a) simplified FE model is pulled in tension with a ramped displacement in the z-direction. The model consists of one cubic hex element and one hundred evenly spaced truss elements. The displacement was applied slowly 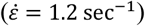, in order to approximate quasi-static loading. The resulting (b) stress response roughly matches the tensile test data [25]. Note that the nominal stress is the current force divided by the original area.

### 2.3 DAMAGE AND HEALING MODELS

#### 2.3.1 STRESS SOFTENING

In previous studies [25] [26], brain tissue has been observed to experience significant stress softening, also called the Mullins effect, after the initial loading cycle. This stress softening behavior has been successfully modeled for brain tissue [25] using the pseudo-elasticity theory [28], which was originally developed for modeling filled rubber elastomers [27]. The softened response curves for the fiber material are shown in Fig. 3. Under a steadily increasing load, the elastic material response follows the primary loading curve. Now, imagine that we have loaded the material along abb’ and stopped at point b’. If we then unload the material from point b’, the response will follow b’Ba, not b’ba. In addition, if we then reload the material from point a, the response will follow aBb’, not abb’. However, after point b’, the response will once again follow the primary loading curve. Therefore, if we reloaded the material from point a to point c’, the response would follow aBb’ and then b’cc’. If we then unloaded the material from point c’, the response would follow c’Ca. Overall, the material softens after the initial loading cycle and the degree of softening depends on the previous strain history. However, the degree of softening is unaffected by the number of loading cycles.

**Fig. 3.**
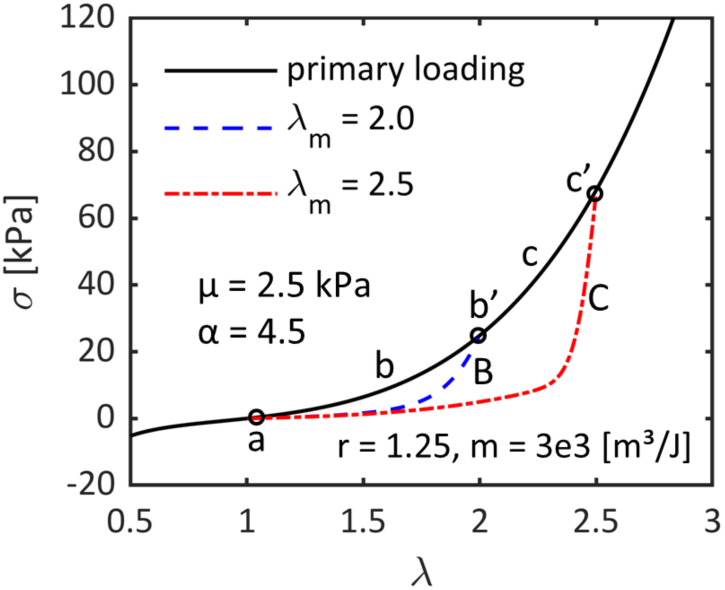
Response curves for the axonal fiber material properties. The dashed (blue) and dash-dot (red) curves represent the softened responses, based on two different levels of previous maximum stretch.

In [28], the pseudo-elasticity theory is presented for an incompressible material experiencing a general biaxial deformation under quasi-static conditions. The governing constitutive relation is:

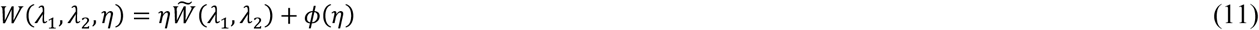

where 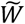 is the undamaged hyperelastic strain energy function, for which the loading and unloading response curves are the same; *η* is a scalar damage variable, which ranges from 0 to 1; and *ϕ*(*η*) is a damage function, which represents the non-recoverable energy associated with the damage process. For the fiber elements, we use Eq. (9) for 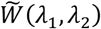. Using Eq. (10) and Eq. (11), the softened uniaxial Cauchy stress is:

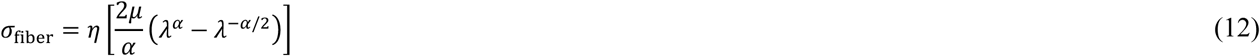

In [28], the scalar damage variable is presented as:

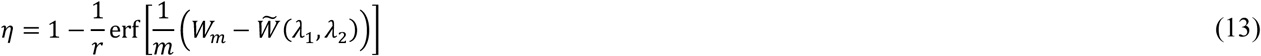

where 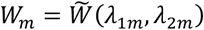; *λ*_1*m*_,*λ*_2*m*_ is the point at which unloading begins (e.g., point b’ or c’ in Fig. 3); *λ*_1_,*λ*_2_ is the current deformation; erf is the error function; and *m* and *r* are positive material parameters. As shown in Eq. (13), *η* is not a fixed scalar value. Instead, it is a function that is used to modulate the stress response in the damaged region (e.g., along path aBb’ or aCc’ in Fig. 3).

In order to implement this stress softening model, we must maintain a record of the minimum stretch *λ_min_* and maximum stretch *λ_max_* experienced by each fiber element, throughout the loading history. When the current stretch is between *λ_min_* and *λ_max_* we calculate the response using Eq. (12). When the current stretch is less than *λ_min_* or greater than *λ_max_* we calculate the response using Eq. (10). In [25], the material constants *m* and *r* were determined for the aggregate brain tissue. However, our composite model requires separate material constants for the fiber and matrix components, which are currently unavailable. After exploring a range of values, as shown in Fig. 4, we chose *m* = 3,000 m^3^/J and *r* = 1.25 as a reasonable qualitative fit.

**Fig. 4.**
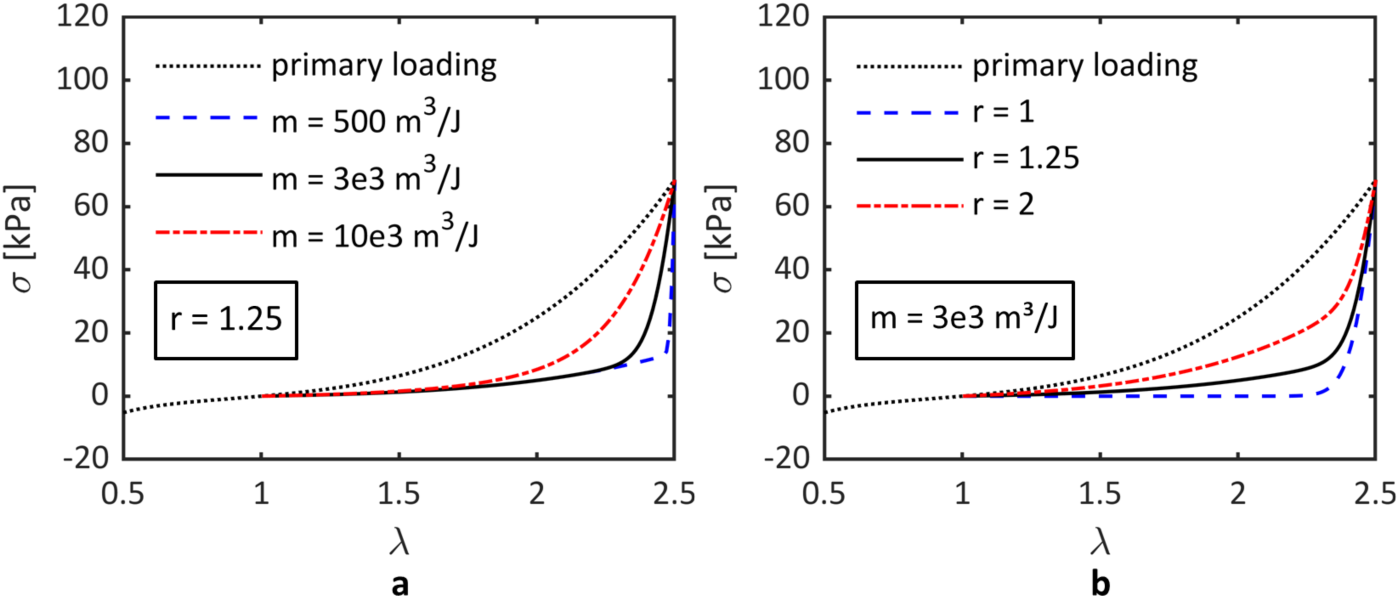
Parametric study used for the selection of material parameters (a) *m* and (b) *r* used in Eq. (13) for the fiber material defined by Eq. (7) and the material properties shown in Table 1. Note that parameter *r* is held constant in (a) and parameter *m* is held constant in (b).

Finally, note that the pseudo-elasticity theory has only been implemented for the fiber component at this time. The fibers are the primary load carrying members. Therefore, we believe this simplification is reasonable for this preliminary work. In the future, we plan to extend the pseudo-elasticity model to the matrix component as well.

#### 2.3.2 FATIGUE SOFTENING

In order to introduce cycle-dependent degradation and finite life, we have also implemented a fatigue-induced stress softening model [30]. At this time, the functional and mechanical fatigue properties of brain tissue have not yet been experimentally studied. However, fatigue behavior has been observed in other soft biological tissues; such as tendons [22], intervertebral discs [23], and bioprosthetic heart valves [24]. Based on the prevalence of fatigue damage in other soft biological tissues, we assume that brain tissue also experiences cycle-dependent degradation.

The following constitutive relation has been used to simulate fatigue damage and permanent set in bioprosthetic heart valves [30]:

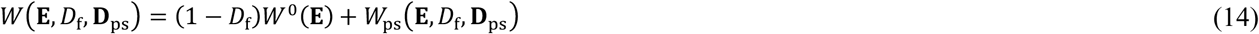

where *W*^0^ is the unfatigued strain energy function; *W*_ps_ is the dissipated strain energy due to permanent set; *D*_f_ is a scalar stress softening parameter associated with fatigue damage, which ranges from 0 to 1; **D**_ps_ is a permanent set parameter; and **E** is the Green strain tensor. The associated PK2 stress tensor is:

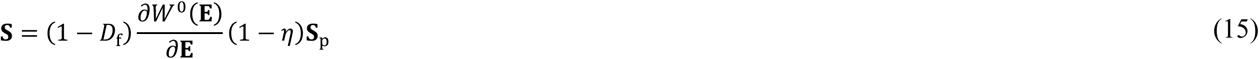

where **S**_p_ is the plastic stress tensor and *η* is a scalar function used to modulate **S**_p_. In part, *η* is a function of the peak strain in the current load cycle. In the referenced study, the authors use a fixed strain amplitude. As a result, the peak strain was constant and known after the initial load cycle. On the of hand, head impacts are unique events. Therefore, the peak strain varies and cannot be determined until each load cycle is complete. In this case, we cannot determine *η* for the current load cycle. Therefore, we cannot properly regulate **S**_p_. As a result, we must ignore permanent set, for this study, and rewrite Eq. (14) in the following simplified form:

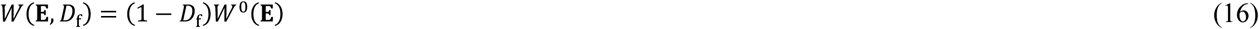

From Eq. (16), the PK2 stress tensor can be expressed as:

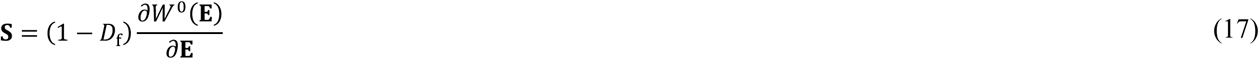

For the fiber elements, we use Eq. (9) for *W*^0^ and Eq. (12) for *σ*_fiber._ Therefore, we can rewrite Eq. (17) as:

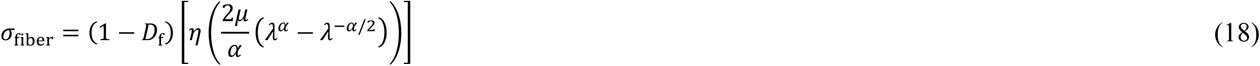

In [30], the increase in *D*_f_ is based on the peak equivalent strain in each load cycle, which is defined as:

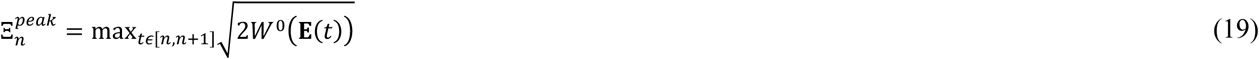

After each load cycle, the number of cycles to failure is calculated for the value of peak equivalent strain using:

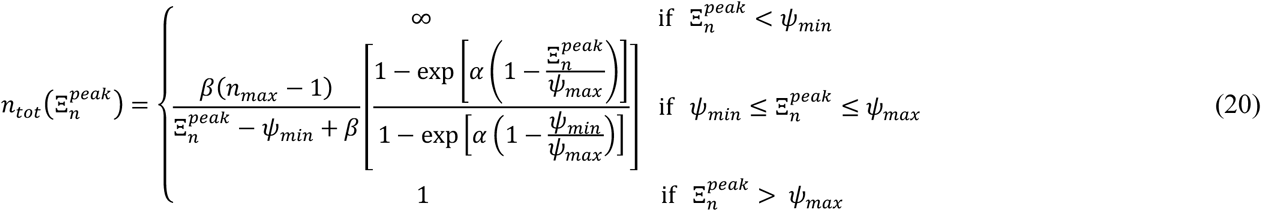

where *ψ_min_* and *ψ_max_* are equivalent strain values that define the boundaries of the fatigue damage zone, *α* and *β* are material properties that control the amount of damage incurred during each cycle, and *n_max_* is the maximum number of cycles permitted at *ψ_min_*. Finally, the stress softening parameter is updated using the following linear damage accumulation model:

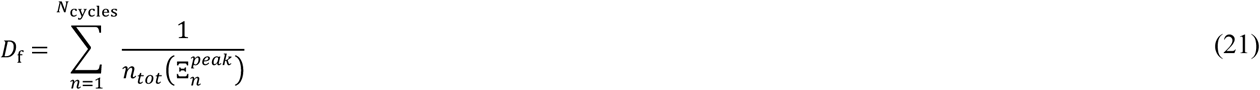

Consider the arbitrary fiber stretch history shown in Fig. 5. In our implementation, we update *D*_f_ after each load cycle is complete, which corresponds to points c, e, and g in Fig. 5. In this case, the stress softening increments in Eq. (21) would be based on the equivalent strain at points b, d, and f. Due to the lack of experimental data, the material properties in Eq. (20) have been estimated based on various practical assumptions. For instance, we chose 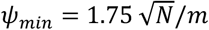, which is based on the assumption that stretch values below 1.02 should not cause fatigue damage. In addition, we assumed that it would be reasonable to sustain a significant amount of cycles at a stretch value of 1.02 before complete mechanical failure. Therefore, we chose *n_max_* = 500. During experimental tension tests on brain tissue [25], final separation occurred at a mean stretch value of 2.66. Therefore, we chose 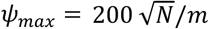, which corresponds to this stretch value. After exploring a range of values, as shown in Fig. 6, we chose *α* = 5 and 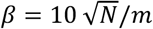 as a reasonable qualitative fit.

**Fig. 5.**
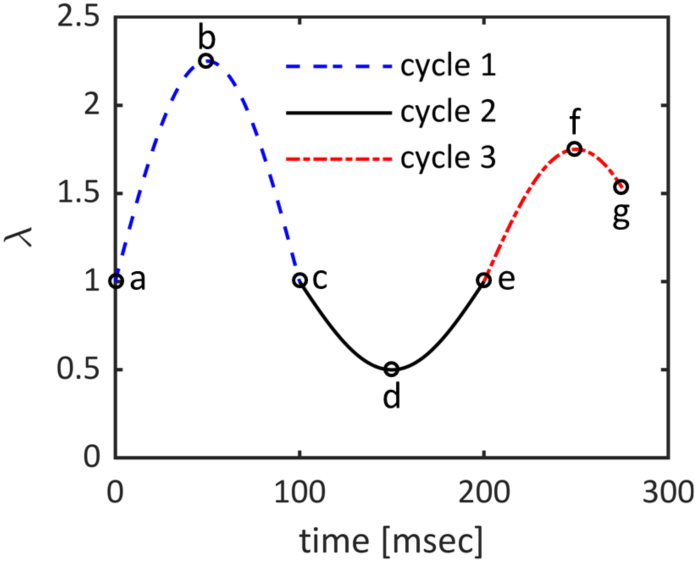
Arbitrary stretch history for a single fiber element. In our implementation, cycles 1 and 2 would be considered full cycles; while cycle 3 would be considered a partial cycle, as it does not return to *λ* = 1. However, note that both full and partial stretch cycles result in fatigue damage.

**Fig. 6.**
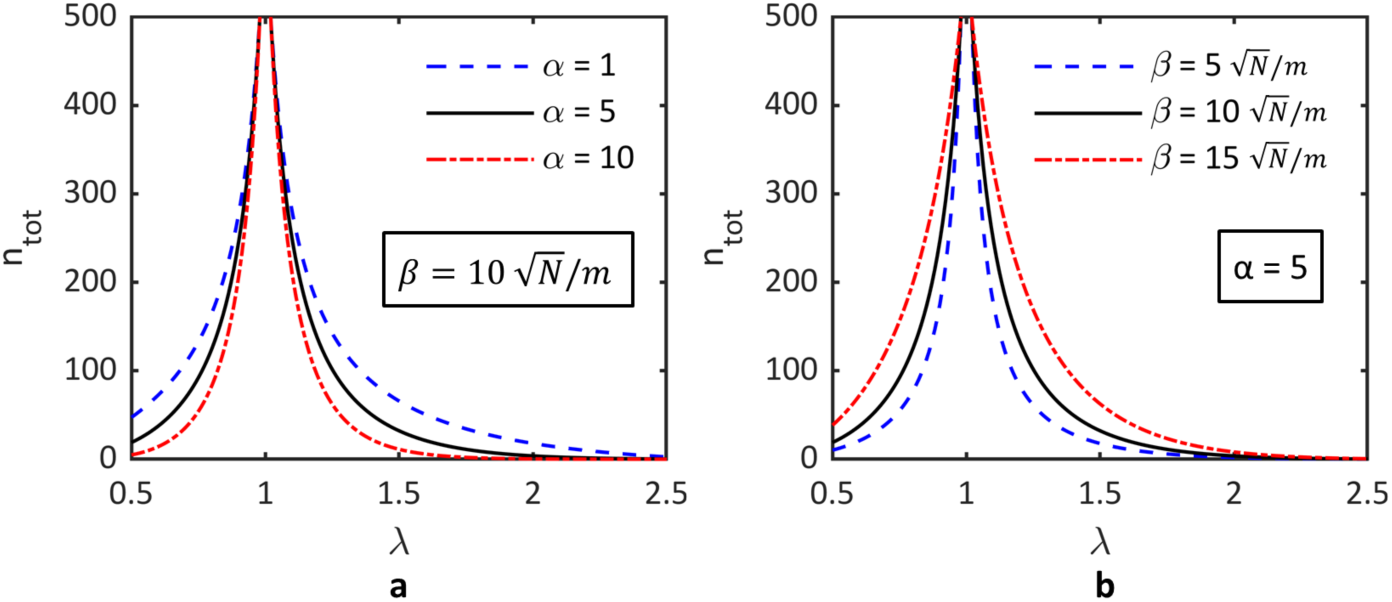
Parametric study used for the selection of the material parameters (a) α and (b) *β* used in Eq. (20) for the fiber material defined by Eq. (7) and the material properties shown in Table 1. Note that parameter *β* is held constant in (a) and parameter α is held constant in (b).

#### 2.3.3 FIBER RUPTURE DAMAGE

Consider the experimental stress response [25] shown in Fig. 2b and observe the softening that takes place after peak stress. Note that a tensile test is a single partial loading cycle, according to Fig. 5. Both the pseudo-elasticity theory and fatigue-induced stress softening models gather information from the initial loading cycle and then alter the response of the second loading cycle accordingly. However, there is no second loading cycle in a tensile test. Therefore, these damage models are not capable of producing the post-yield softening behavior seen in Fig. 2b. In order to address this shortcoming, we have included an additional damage parameter, which also ranges from 0 to 1. The softening that occurs before failure is typical of damaging fibrous materials and is caused by the progressive rupture of the filamentary structure [25]. Therefore, the additional damage parameter is referred to as fiber rupture damage, *D*_r_. Based multiple quasi-static tensile tests [25], fiber damage initiates at a mean stretch of 1.91 and final separation occurs at a mean stretch of 2.66. As a preliminary attempt, we have assumed a linear evolution of fiber rupture damage, based on the fiber stretch history:

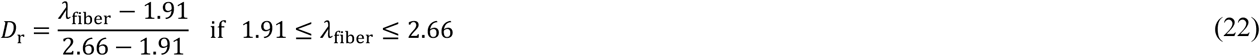

The total fiber damage, *D*_tot_, which ranges from 0 to 1, is then calculated for each fiber as the sum of the fatigue and fiber rupture damage parameters:

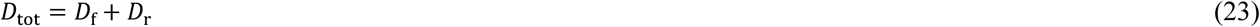

Finally, using Eq. (23), we can rewrite Eq. (18) as:

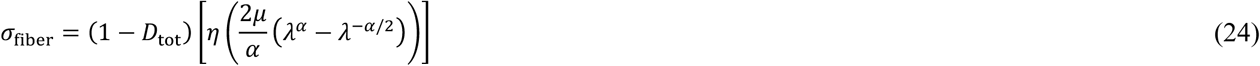

#### 2.3.4 MATRIX DAMAGE

So far, we have only considered fiber damage. In order to incorporate matrix damage, we first calculate the average fiber damage for the fibers associated with each matrix element as follows:

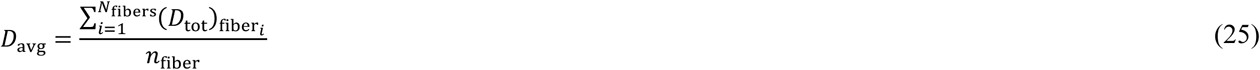

Then, using Eq. (17), we can calculate the damaged matrix stress as follows:

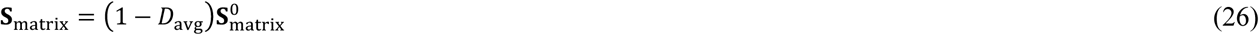

where 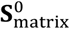 is the undamaged matrix PK2 stress tensor.

#### 2.3.5 HEALING

At the end of each simulation, we export *λ*_min_, *λ*_max_, *D*_f_, and *D*_r_ for each fiber element. We then import these scalar damage parameters as the initial conditions for the next impact simulation. In addition, we have provided a mechanism to account for the healing process between two successive impacts. When the next simulation is initialized, a healing duration can be specified, which then partially or completely reverses the accumulated damage in each fiber element. Healing models have been implemented in various simulations of soft biological tissues, such as tendons [31], ligaments [32], and intervertebral discs [33]. However, due to the lack of experimental data, a constant healing rate is typically assumed. Similarly, we also assume a constant healing rate. In addition, we have assumed exponential recovery functions, which are defined as follows:

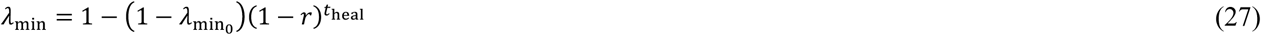

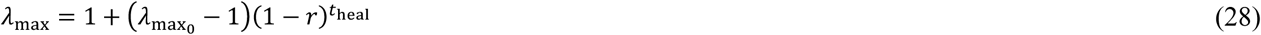

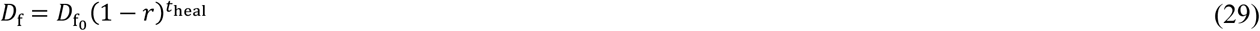

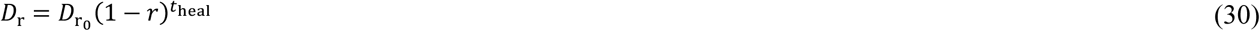

where 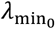, 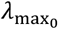, 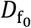, 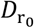 are the initial values of the damage parameters exported from the previous simulation; *r* is the constant healing rate, where we have assumed *r* = 0.025; and *t*_heal_ is the healing duration in number of days. For reference, Eq. (29) has been plotted in Fig. 7 for various values of *r*. The plots for Eq. (27) – Eq. (30) are similar. However, note that *D*_f_ and *D*_r_ recover to zero, while *λ*_min_ and *λ*_max_ recover to 1.

**Fig. 7.**
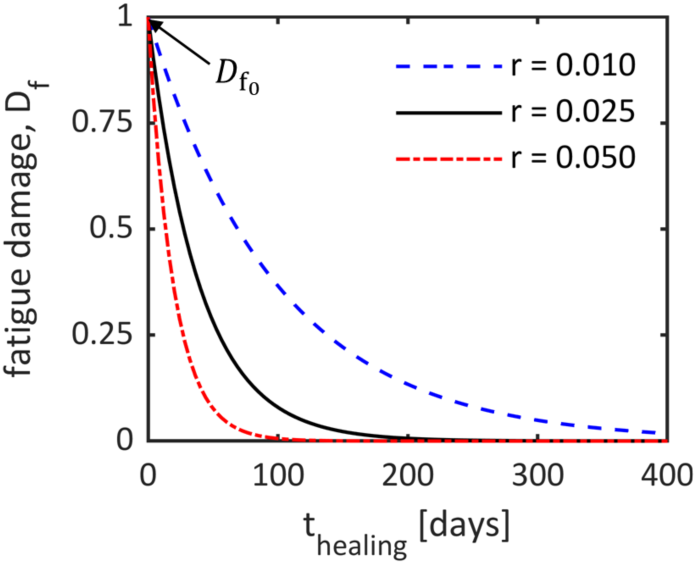
Plot of Eq. (29) using an arbitrary value of initial fatigue damage, 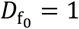. The dashed (blue) and dash-dot (red) curves illustrate how alternative values of *r* affect the recovery of the damage parameter.

Many researchers have studied the recovery process that follows TBI [39] [40] [41] [42] [43]. However, a definitive relationship between recovery time and impact severity, location, and frequency has not yet been established. Most studies provide trends associated with a specific injury type and setting. Therefore, it is difficult to combine these separate findings into a single comprehensive formula. For example, in the case of minor sports injuries, symptoms tend to subside in seven to ten days [44]. However, for more severe injuries, many TBI patients go on to develop post-concussive syndrome (PCS), which is associated with long term cognitive deficits and white matter changes [45] [46] [43]. Due to the lack of a generalized recovery criteria, we have assumed a simple exponential healing model that provides a full recovery after six months (i.e., *r* = 0.025), which is based on a three to six month recovery time for the typical civilian mTBI [45].

Finally, it is important to recognize that healing is an extremely complex remodeling process that involves many interacting biochemical and biomechanical processes [32]. In addition, the repair process can often lead to the formation of new structures, such as glial scars, which results in altered mechanical properties [4]. In addition, it is important to note that damage modeling is concerned with mechanical behavior and structural integrity, while most recovery studies are concerned with patient symptoms and functional restoration. Therefore, in order to accurately model damage recovery, future research is needed to understand how mechanical damage recovers over time.

### 2.4 FE MODEL PREPARATION

The FE model is based on diffusion tensor imaging (DTI) scans of a male rugby player, which were obtained from the Pennsylvania State University Center for Sports Concussion Research and Service. The scans were performed in a Siemens Trio Tim 3.0T MRI machine. We used Diffusion Toolkit [47] to generate the fiber track data and TrackVis [47] to visualize the fiber tracks, as shown in Fig. 8a. In addition, we used a custom MATLAB script to convert the binary TrackVis track file to an ASCII mesh file, which consists of nodal coordinates and element connectivity. We then used another custom MATLAB script to extract a cubic sample from the full tractography, as shown in the boxed region of Fig. 8b. We purposely selected a region of complex fiber geometry, which includes fiber crossings and branching, in order to demonstrate the robustness of the embedded element method. The final FE model, as shown in Fig. 8c, contains 2,994 individual fiber elements, with a mean length of 3.4 mm. Note that a single fiber track consists of multiple fiber elements. For example, in Fig. 8a, the mean fiber track length is 22.5 mm.

**Fig. 8.**
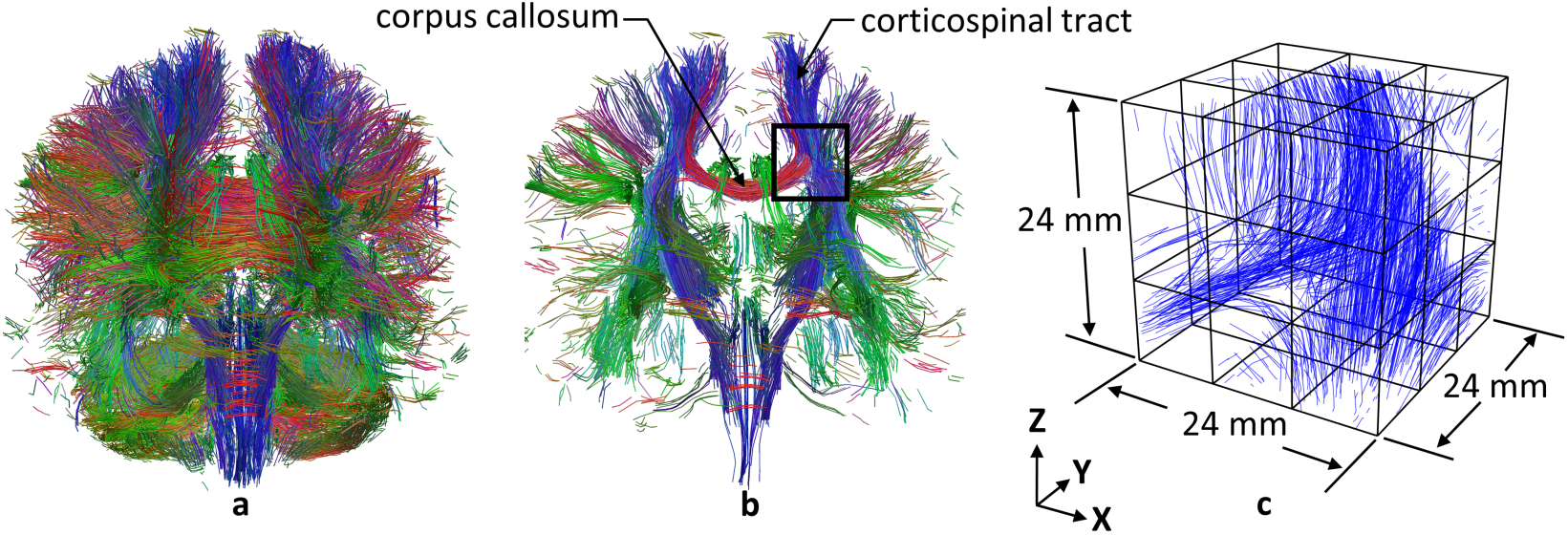
Tractography for (a) the full brain and (b) a coronal cross-section of the full brain. The (c) current FE model is based on the boxed region in (b).

In order to study the evolution of damage in the fiber elements and the resulting effect on the global tissue response, we prepared multiple example simulations using various simple loading configurations. For instance, consider Fig. 9a, where we have loaded a single face in the z-direction. In this case, we provide minimal restraints by fixing the nodal degrees of freedom shown in Fig. 9b. The resulting model behavior is illustrated in Fig. 10. Note that if we load a single face in the x-direction, as shown in Fig. 9c, the restraints will be similar to those shown in Fig. 9b, but not identical.

**Fig. 9.**
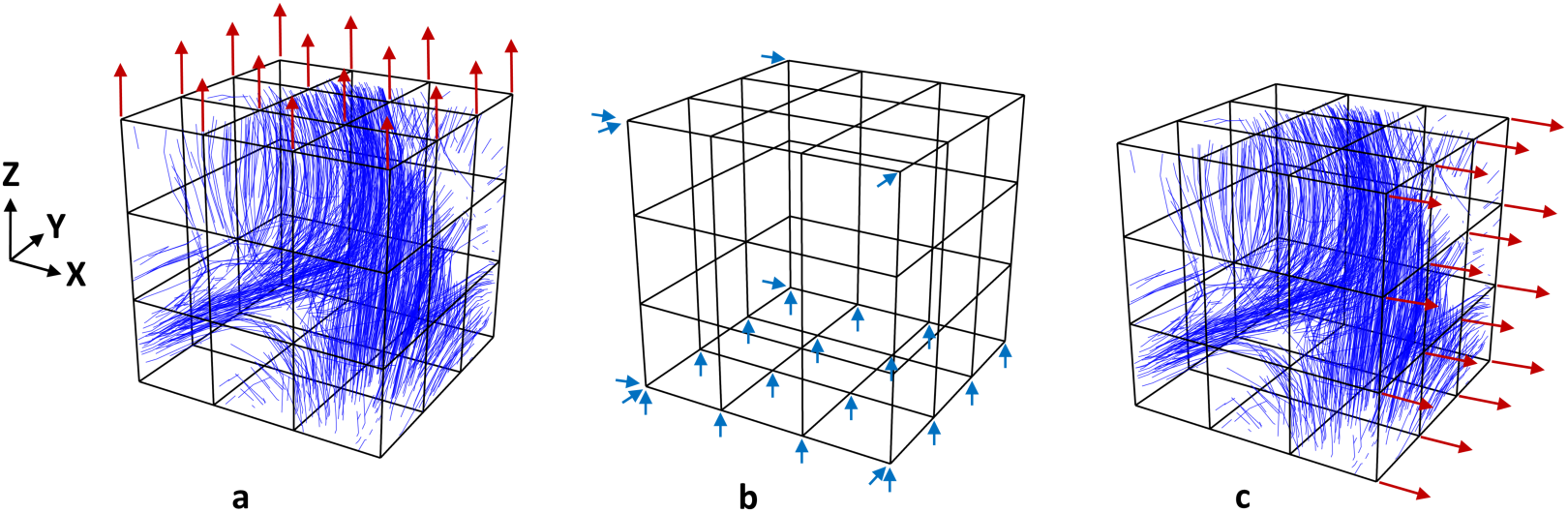
Applied load in the (a) z-direction and the associated (b) nodal restraints. The arrows in (b) indicate the nodal displacement components which are fixed, for the z-direction loading. An applied load in the (c) x-direction will require similar restraints, but not identical to those shown in (b).

**Fig. 10.**
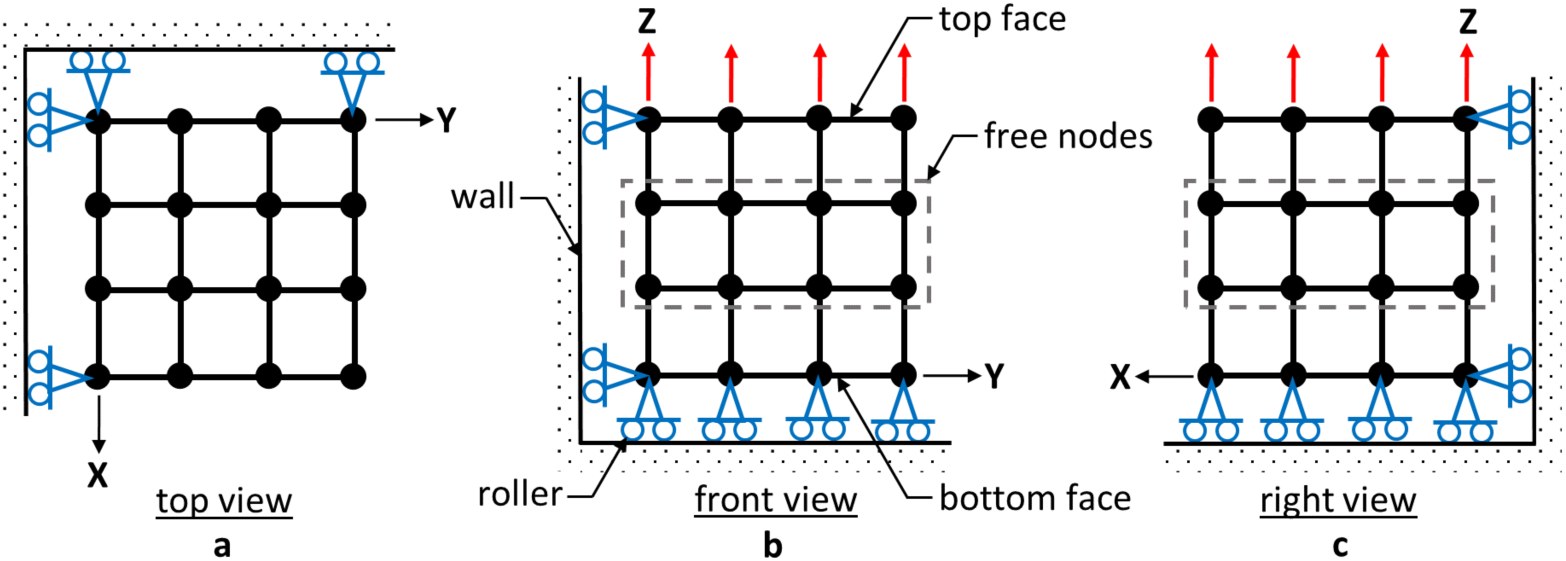
Behavior of the FE model when subjected to the applied load shown in Fig. 9a and the nodal restraints shown in Fig. 9b. Note that the rollers remain in contact with the wall at all times. Also, note that the middle region of the FE model is unrestrained and free to move in any direction. This allows us to visualize the irregular distortions caused by the nonuniform distribution of the fiber elements.

To select an appropriate matrix element size, we completed a mesh convergence study. Specifically, we applied a ramped 16 mm displacement in the z-direction and studied the effects of matrix element size on the stress response of a single fiber element. Note that the 16 mm displacement magnitude is arbitrary and was selected only to produce a significant amount of model deformation. The ramped load profile is shown in Fig. 11 and the three test models are shown in Fig. 13b. For this study, we selected the arbitrary fiber element (i.e., truss element 1082) shown in Fig. 12, due to its close alignment with the direction of loading. As seen in Fig. 13a, the fiber stress is not very sensitive to matrix element size. The final stress in the 3 × 3 × 3 configuration is only 1.6% smaller than the final stress in the 5×5×5 configuration. However, the 5×5×5 configuration simulation time was 2.1 times longer. Therefore, we choose the 3×3×3 configuration for our example simulations, which provided a reasonable balance of accuracy and simulation time. The 3×3×3 configuration consists of twenty-seven 8 mm cubic hex elements.

**Fig. 11.**
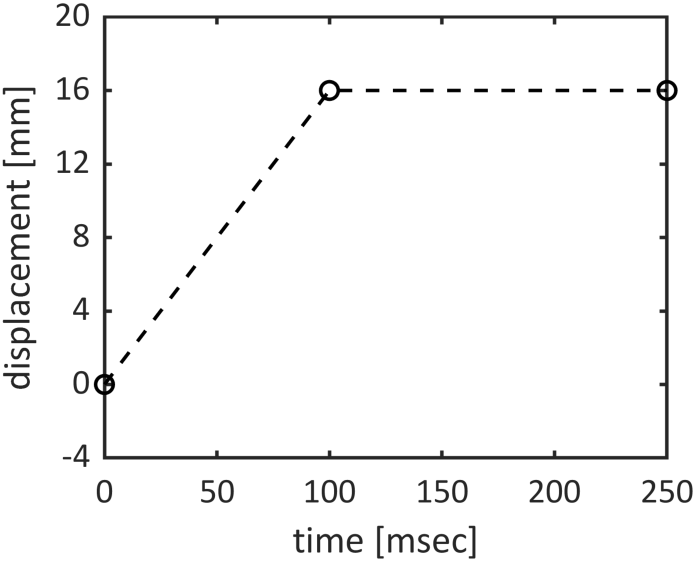
Ramped load profile.

**Fig. 12.**
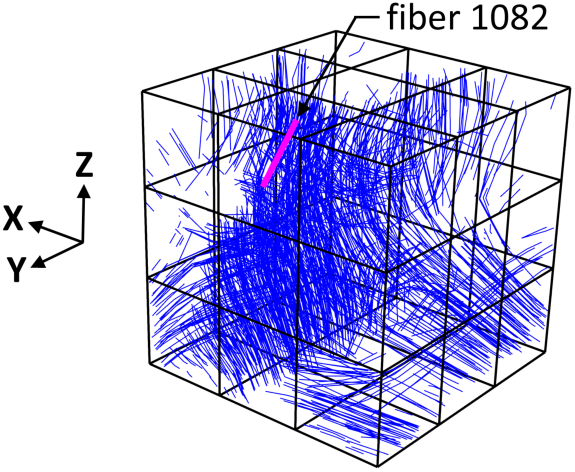
An arbitrary fiber element selected for close examination.

**Fig. 13.**
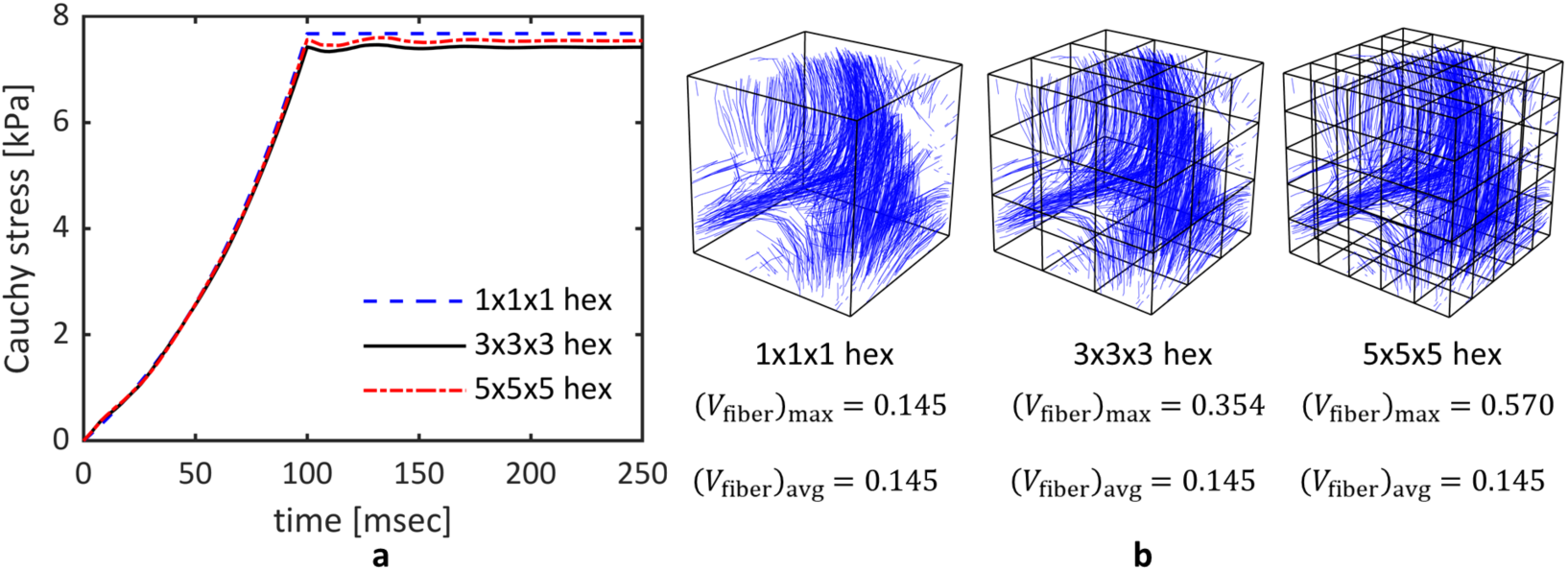
The (a) results of the mesh convergence study and the (b) associated mesh options. In all three cases, a ramped 16 mm displacement was applied in the z-direction. The fiber stress in (a) is associated with the fiber element highlighted in Fig. 12. Note that all damage features were disabled for this mesh convergence study.

Finally, we needed to specify a cross-sectional area for each truss element. A fiber-reinforced composite contains a mixture of fiber and matrix components. Depending on the shape and packing of the fibers, there are limits on the fiber volume fraction, which is defined as:

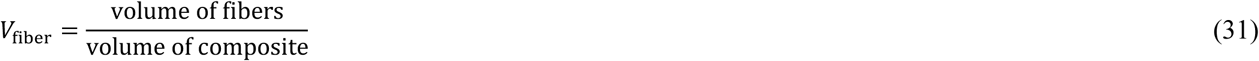

For the ideal case of round parallel fibers, the maximum fiber volume fraction is 0.785 for square packing and 0.907 for hexagonal packing [35]. Axonal fiber tracts are not typically arranged in an ideal parallel arrangement. Therefore, we chose 0.785 as reasonable maximum for the fiber volume fraction in a single matrix element. By trial-and-error, we discovered that a fiber element diameter of 0.5 mm (area = 1.9635e-7 m^2^) produced the maximum allowable fiber volume fraction in the 5×5×5 hex configuration, shown in Fig. 13b. For reference, the maximum and average fiber volume fractions for all three mesh configurations are provided in Fig. 13b, for a fiber element diameter of 0.5 mm.

### 2.5 SIMULATION OVERVIEW

In order to demonstrate the features and mechanics of the proposed computational framework, we present two example simulations. In the first example, we highlight a gap in the current technology and show how damage modeling offers a potential solution. In the second example, we step through the complete workflow of a practical repetitive impact problem. Along the way, we review the evolution of damage and its effect on the aggregate tissue behavior. In both examples, we use unrealistically large displacements, in order to produce a significant amount of damage in a small number of cycles. Through trial and error, we discovered that a 16 mm displacement magnitude was sufficient to clearly illustrate the damage behavior. Otherwise, the 16 mm value is arbitrary and has no physiological significance. For reference, a 16 mm displacement is equivalent to a nominal stretch of 1.67, for the current 24 mm cubic FE model.

## 3 RESULTS

### 3.1 EXAMPLE 1 – RAMPED LOAD

In this first example, we begin with a demonstration of the current technology and its limitations. Therefore, the damage features are initially disabled in the code. We start off with the FE model shown in Fig. 8c. Then, we apply a ramped 16 mm displacement, per Fig. 11, in the z-direction, as shown in Fig. 9a. The resulting fiber strains are presented in Fig. 14a. Next, we complete a separate simulation with a ramped 16 mm displacement in the x-direction, as shown in Fig. 9c. The resulting fiber strains are presented in Fig. 14b. Simulations of this type are useful because they allow us to compare the resulting fiber strains with established tissue thresholds, such as those presented in [21]. However, each simulation represents a separate isolated loading event. Currently, there is no way combine the results of successive head impact simulations. Therefore, it is impossible to monitor the long-term structural health of brain tissue. In addition, the tissue does not degrade. Therefore, the material behavior remains constant throughout all simulations. These are significant limitations with the current technology. However, damage modeling can be used to address these issues.

**Fig. 14.**
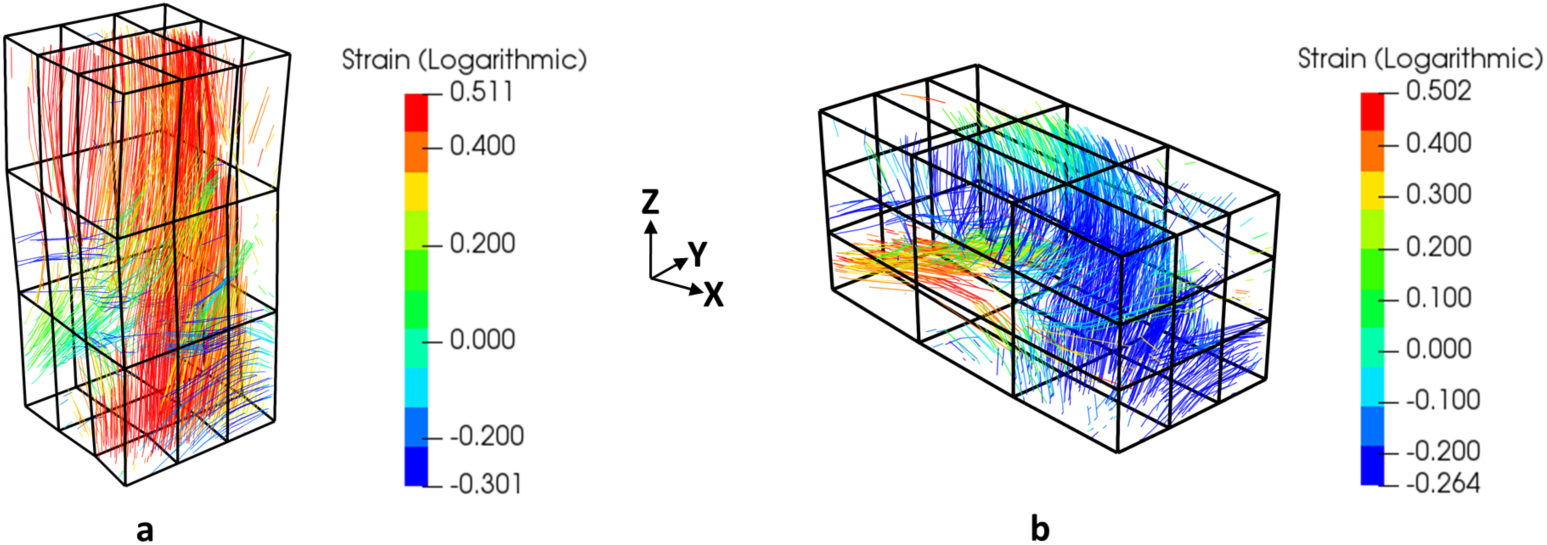
Fiber strain results for a single ramped 16 mm displacement applied separately in the (a) z-direction and the (b) x-direction. Note that these are two completely independent simulations and all damage features have been disabled.

We now reconsider the previous two simulations. This time, we enable all damage features in the code. First, we simulate the applied displacement in the z-direction. Now, we plot the resulting fiber damage *D*_tot_ in Fig. 15a. Note that we are below the stretch threshold for fiber rupture damage to occur. Next, we import the damage parameters *λ*_min_, *λ*_max_, *D*_s_, and *D*_r_ into the second simulation and apply the displacement in the x-direction. The resulting fiber damage plot is shown in Fig. 15b. By introducing damage, we are now able to view the consequences of successive impacts. In Fig. 15b, we see that many fibers have suffered a significant level of damage. However, the fibers inline with and perpendicular to the x and z axes have suffered the most damage, while the diagonal fibers have suffered the least damage. This result can be explained using the following simple example. Consider an arbitrary incompressible (i.e., constant area) unit square, as shown in Fig. 16a. Now, imagine that we apply an arbitrary stretch of 2.00 in the x-direction, as shown in Fig. 16b. The resulting lateral stretch (i.e., y-direction) would be 0.50 and the resulting diagonal stretch would be 1.46. Using Eq. (20) and Eq. (21), we can calculate the fatigue damage associated with each of these three stretch values. For the stretch values of 2.00, 0.50, and 1.46, the resulting fatigue damage values are 0.269, 0.053, and 0.026, respectively. Based on this example, the results in Fig. 15 are now somewhat intuitive. However, if we were to use a more complicated sequence of applied loads, the combined fiber damage results would not be as obvious.

**Fig. 15.**
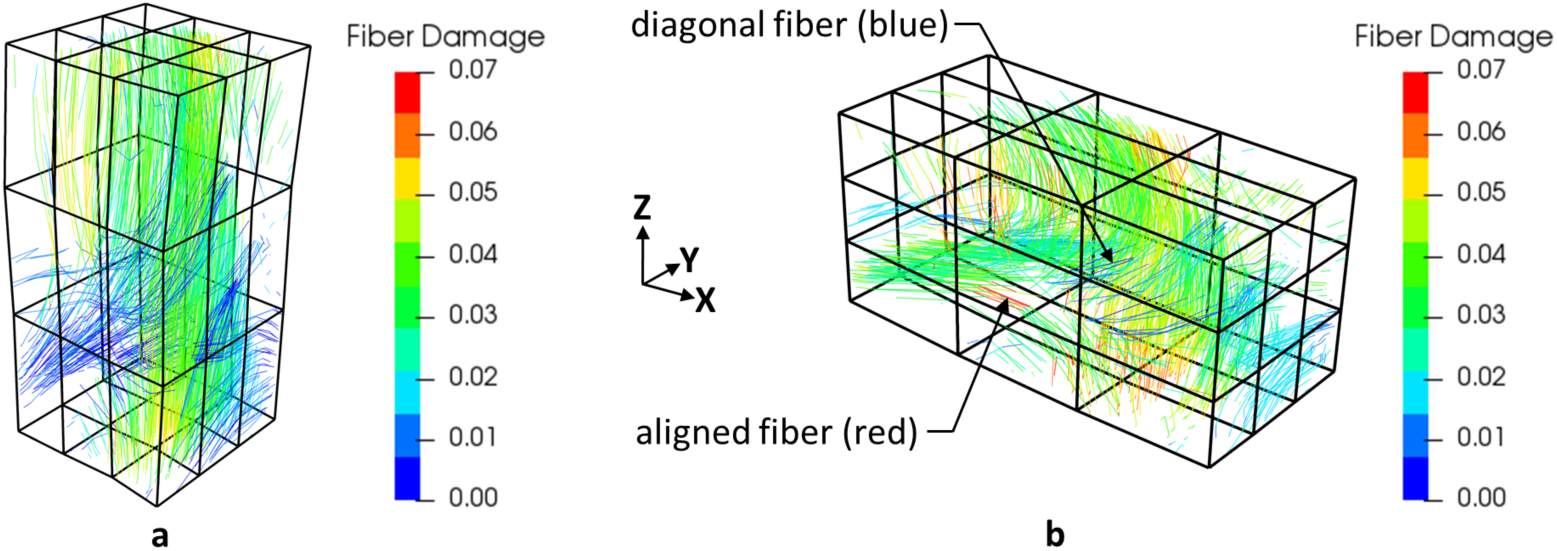
Fiber damage results for a single ramped 16 mm displacement applied consecutively in the (a) z-direction and then the (b) x-direction. The fiber damage results from (a) were imported as the initial conditions for (b). Fibers in-line with and perpendicular to the loading directions suffer the most damage, while diagonal fibers suffer the least damage.

**Fig. 16.**
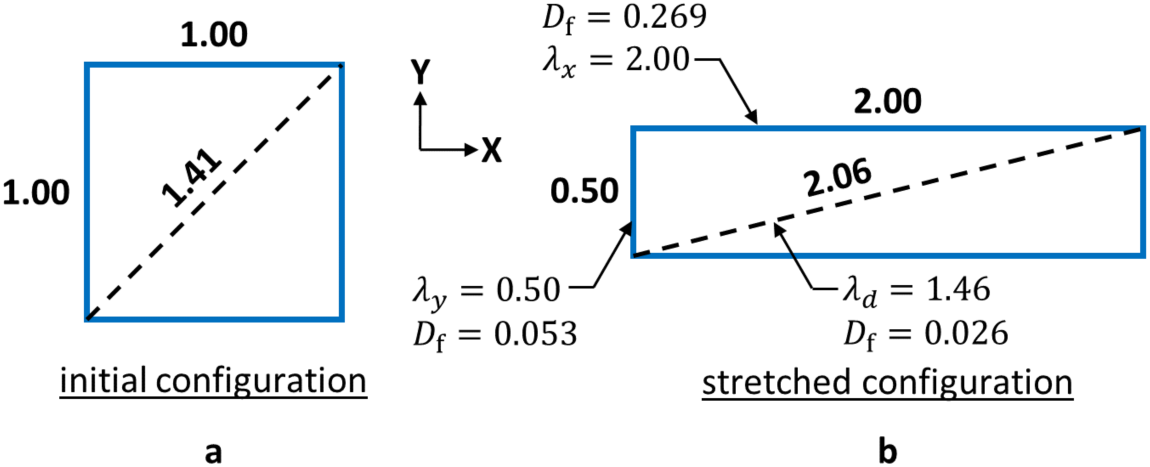
An incompressible (i.e., constant area) unit square shown (a) before deformation and (b) after deformation. The resulting stretch and fatigue damage values are indicated.

### 3.2 EXAMPLE 2 – CYCLIC LOAD

In this second example, we study a slightly more complex scenario and explore the damage response in greater detail. Again, we start with the FE model shown in Fig. 8c. However, we now apply a cyclic displacement, per Fig. 17, in the z-direction. Injurious head impacts are typically associated with interaction times of 15 msec or less [48]. However, in this example, we applied the load over 100 msec duration ramps, in order to minimize the solution noise caused by rapid changes in load profile. One should note that such solution noise is normal and also observed when using commercial software.

**Fig. 17.**
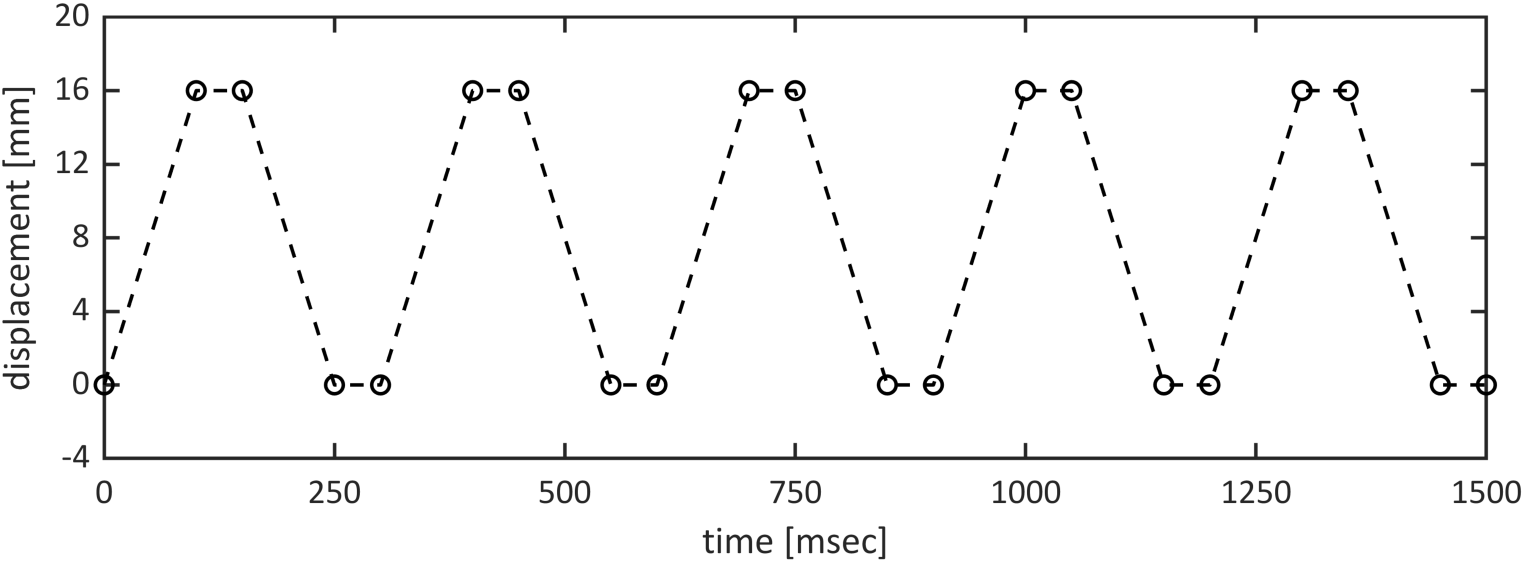
Cyclic load profile.

In Fig. 18, we plot the stress response for the fiber element shown in Fig. 12. During the first load cycle, the effects of damage are minor. However, we see the response change shape on the unloading portion of the cycle, due to stress softening (i.e., the Mullins effect), per Eq. (12). This change in shape is then present in each of the following load cycles. At the end of each cycle, the additional fatigue damage is calculated using Eq. (21). As a result, we see the stress amplitude drop after each load cycle, according to Eq. (18). As previously mentioned, the nominal stretch is 1.67 in these examples. Therefore, fiber rupture damage does not occur, according to Eq. (22). At the peak of the fifth load cycle (1325 msec), the fiber stress has decreased by 17.5%. The final total fiber damage is 0.22.

**Fig. 18.**
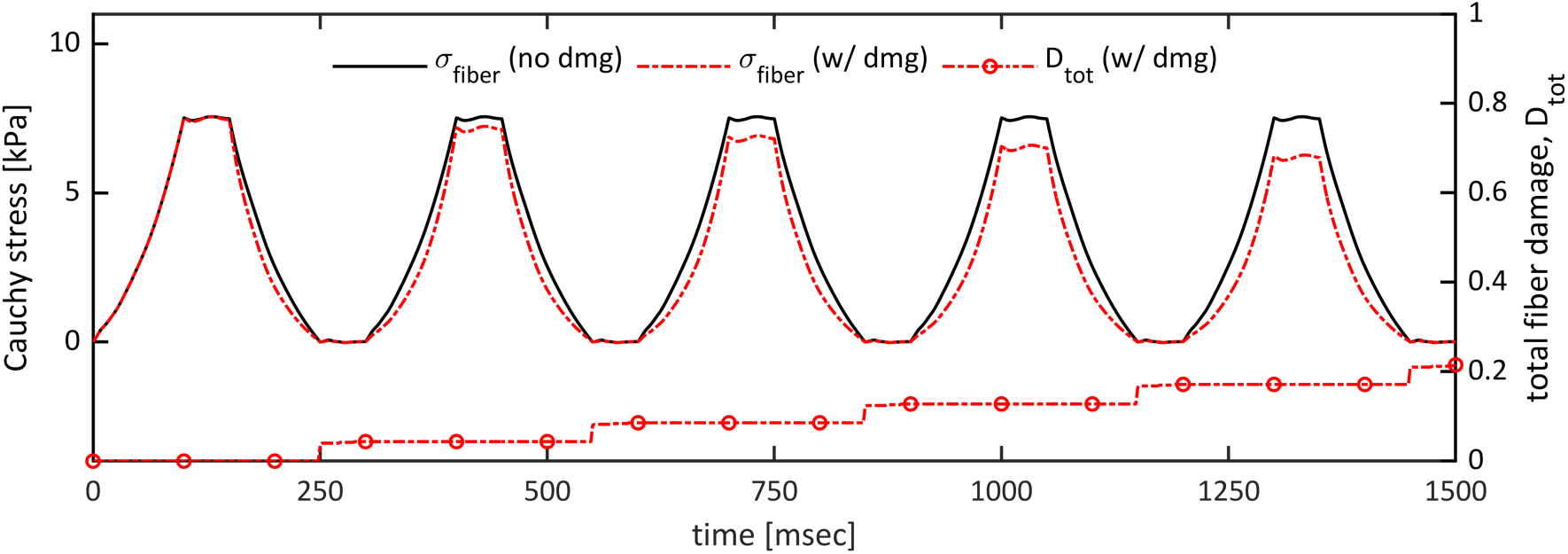
The stress response of fiber element 1082 during the initial five loading cycles on day 1. The total fiber damage for fiber element 1082 is also shown. Note that the word “damage” has been abbreviated as “dmg.”

Now, imagine that some time has passed. For the sake of this example, assume that it is three months (i.e., 91 days) later. We now repeat the previous five displacement cycles. However, in this case we will consider two options. In the first option, we allow damage to progress as before. This is equivalent to ten successive load cycles. In other words, the three-month rest period has no effect on the results. In the second option, we use Eq. (27) – Eq. (30) to partially restore the damage parameters before the initial load cycle begins.

In Fig. 19, we plot the fiber stress response for both options (i.e., healing vs. no healing), for the fiber element shown in Fig. 12. At the peak of the first load cycle (125 msec), the unhealed fiber stress is 21.9% less than the undamaged response, whereas the healed fiber stress is 25.4% greater than the unhealed fiber stress. At the peak of the fifth load cycle (1325 msec), the unhealed fiber stress is 39.4% less than the undamaged fiber stress, whereas the healed fiber stress is 32.8% greater than the unhealed fiber stress. For the unhealed option, the total fiber damage was initialized at 0.22 and finished at 0.43. For the healed option, the total fiber damage was initialized at 0.02 and finished at 0.24.

**Fig. 19.**
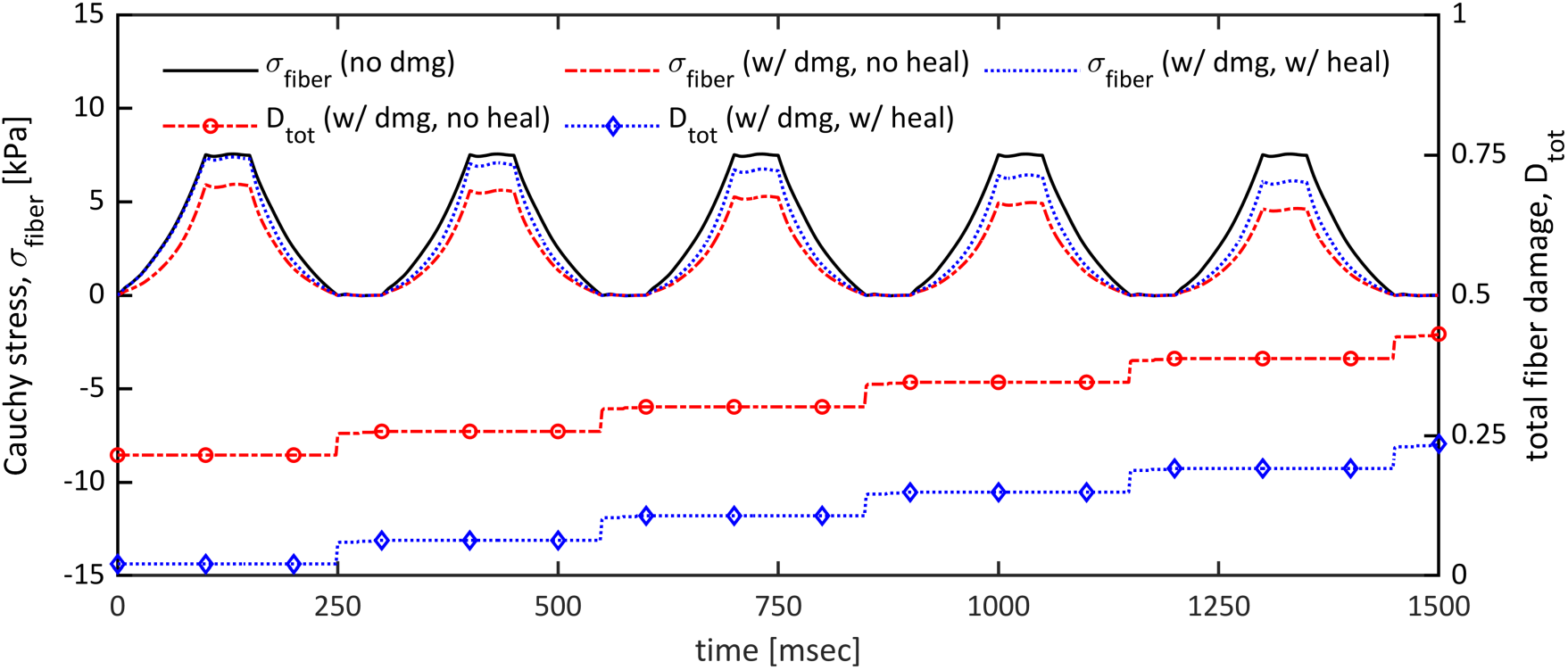
The stress response of fiber element 1082 during the second five loading cycles on day 91. The total fiber damage for element 1082 is also shown. Note that the word “damage” has been abbreviated as “dmg.”

## 4 DISCUSSION

Current FE models are unable to capture the progressive damage caused by repeated head trauma. In this study, we present a method for computing the history-dependent mechanical damage of axonal fiber tracts in the brain. The proposed fiber damage model consists of three main components. First, we use the pseudo-elasticity theory to account for stress softening (i.e., the Mullins effect). Next, we use a fatigue-induced stress softening model to provide cycle-dependent degradation and finite life. Lastly, we use a fiber rupture damage model to generate post-yield softening behavior. In addition, we use an exponential healing model to partially or completely reverse the accumulated damage in each fiber element. Using two examples, we have demonstrated the ability of the proposed model to successfully track cumulative axonal damage and degrade the mechanical response of the aggregate brain tissue.

At this time, the greatest barrier to further development is the lack of experimental data. Stress softening has been observed in human brain tissue [25]. However, the fatigue properties of brain tissue have not yet been studied. Therefore, we relied on parametric studies and our best judgment to select reasonable values for the required model parameters. Due to the soft and fragile nature of brain tissue, it is challenging to perform mechanical tests [26]. However, it is even more difficult to study the effects of healing on the mechanical properties of living brain tissue. As a result, such data does not currently exist. Many studies have looked at the recovery of TBI symptoms over time. However, it is not clear how the recovery of mechanical properties and the recovery of TBI symptoms are related. Therefore, as with damage, the lack of experimental data is the greatest limitation for modeling the recovery process.

The embedded element method is advantageous because it preserves the complete fiber track resolution provided by the tracking software. This makes it relatively simple to identify potentially damaged fiber tracts, based on the simulation results. However, the embedded element method requires separate material properties for the matrix and fiber components, which exacerbates the problem of limited experimental data. In the end, we may need separate elastic, damage, and recovery properties for the matrix and fiber components. Also, experimental data is typically supplied at the composite tissue level. Therefore, it is necessary to decompose the results into component level properties.

In this study, we have focused on mechanical damage. However, we are also interested in predicting functional impairment. There is likely some connection between mechanical damage and functional injury. However, it is unlikely that there is a one-to-one relationship. Tension tests [25] have shown that brain tissue supports mechanical load up until fracture, at a mean stretch value of 2.66. On the other hand, electrophysiological impairment been shown [21] to occur much earlier, at a stretch value of 1.17. Therefore, as more experimental data becomes available, it may become necessary to introduce a separate set of functional damage parameters that will be tracked in parallel with the proposed mechanical damage parameters.

In its current form, the proposed computational framework should be regarded as a first step in a necessary direction. To be useful in practice, the proposed models must be refined and validated. Nonetheless, the current model has immediate value in that it provides a new metric (i.e., inelastic damage) that captures both the magnitude and frequency of successive fiber tract strains; a capability which otherwise does not exist. In addition, the proposed model also accounts for the changes in the material behavior that result from fiber damage. This new feature has important consequences. For instance, consider an applied cyclic load with a constant force magnitude. As damage accumulates with each additional cycle, the resulting tissue deformation will steady increase. As a result, damaged tissue will be more susceptible to strain-related functional impairment.

Overall, there is a need for long-term brain health monitoring technology. After an initial TBI, the brain enters a temporary period of increased susceptibility to further TBI and long-term deficits. This vulnerable cerebral state is known as the “window of vulnerability” [46]. As a result, it is crucial to identify and treat TBI early on, in order to avoid more severe and permanent injuries from developing [45]. Microstructural white matter damage is thought to be associated with multiple forms of brain injury; such as PCS [46] [49] and chronic traumatic encephalopathy (CTE) [50]. Therefore, we believe that it is especially useful to quantify the potential cumulative damage of axonal fiber tracts in the brain, which result from repeated TBI.

## 5 CONCLUSION

In this paper, we proposed a method for computing the history-dependent mechanical damage of axonal fiber tracts in the brain. Using two detailed examples, we demonstrated the ability of the proposed model to track cumulative damage and degrade the mechanical response of the material. As an additional benefit, fiber damage provides a new metric that captures both the magnitude and frequency of successive fiber tract strains; a capability which otherwise does not exist.

In the future, we plan to extend the current study to the full brain. In addition, we plan to research more sophisticated methods of calculating fiber rupture damage and matrix damage. We would also like to further explore the available options for incorporating permanent deformation, which has been observed in brain tissue [25]. Most importantly, we hope to calibrate and refine the proposed model, when new experimental data becomes available. Based on this initial study, damage modeling has the potential enhance current brain modeling techniques.

## ACKNOWLEDGEMENTS

The authors gratefully acknowledge the support provided by the U.S. Army Research Laboratory (ARL) at Aberdeen Proving Grounds under contracts W15P7T-10-D-D416 and DOTC-17-01-INIT0086, and CFD Research Corporation under a subcontract funded by the Department of Defense, Department of Health Program through contract W81XWH-17-C-0216. All imaging data used here are provided by The Pennsylvania State University Center for Sports Concussion Research and Service, University Park, USA. The authors thank Dr. Sam Slobounov and Dr. Brian D. Johnson for the data provided. The authors also thank Dr. Daniel Cortes at The Pennsylvania State University for his helpful feedback and guidance. Finally, the authors acknowledge Ritika Menghani and Srikumar Sridhar for their significant development contributions to the custom FE code used in this study.

